# Thermodynamics of consciousness: A non-invasive perturbational framework

**DOI:** 10.64898/2025.12.09.691422

**Authors:** Tomas Berjaga-Buisan, Juan Manuel Monti, Martina Cortada, Michele A. Colombo, Sebastian M. Geli, Gianluca Gaglioti, Simone Sarasso, Morten L. Kringelbach, Maurizio Corbetta, Maria V. Sanchez-Vives, Marcello Massimini, Yonatan Sanz Perl, Gustavo Deco

## Abstract

The quest for reliable and objective measures of consciousness is critical in basic and clinical neuroscience. Across species, the Perturbational Complexity Index (PCI) has emerged as a robust empirical marker by directly perturbing the brain, yet its underlying principles of physics are not fully understood. Here, we bridge this gap by introducing a non-invasive framework based on generative whole-brain models of non-equilibrium brain dynamics. Using these models, we identified violations of the Fluctuation-Dissipation Theorem (FDT) in humans and rodents across wakefulness, anesthesia, and disorders of consciousness. Mirroring the patterns observed with PCI, we found decreased FDT violations in unresponsive disorders of consciousness and anesthesia compared to conscious conditions. This reveals a close link between PCI and non-equilibrium dynamics in spontaneous brain signals, grounding PCI in fundamental principles of physics. Overall, this framework offers new complementary, non-invasive, model-based avenues for understanding the nature of consciousness and for developing objective tools to assess its loss and recovery in health and disease. It also provides a principled foundation for discovering novel strategies to restore consciousness.

## INTRODUCTION

Distinguishing between conscious and unconscious brain states remains one of the most fundamental and unresolved challenges in systems and clinical neuroscience. Clinical diagnosis currently relies on a patient’s ability to interact with the environment, which is typically assessed using standardized behavioral scales, such as the Coma Recovery Scale–Revised (CRS-R) ^1,2^. However, the absence of behavioral signs cannot be considered definitive evidence of unconsciousness, as individuals may be unable to interact with the environment due to impairments in motor, sensory, or executive functions ^3–5^. States of disconnected consciousness can occur in healthy individuals during dreaming, under general anesthesia, and in cases of severe brain injury ^6–9^. Therefore, there is an urgent need for objective neurophysiological markers capable of indexing consciousness independently of behavior.

The Perturbational Complexity Index (PCI) is a generalizable non-behavioral marker of consciousness which quantifies the algorithmic complexity of spatiotemporal brain patterns in response to external perturbations, typically delivered via transcranial magnetic stimulation (TMS) in humans or intracortical electrical stimulation in animal models ^10,11^. PCI captures the presence of both functional integration and differentiation, by computing the extent and information content of the chain of interactions evoked in the brain by a direct perturbation. These features are collectively referred to as brain complexity, which has been proposed as a fundamental requirement for consciousness in both theoretical models and neuroimaging studies ^12–14^. PCI has demonstrated reliability in discriminating consciousness from unconsciousness ^15^. Notably, previous studies reported that PCI generalized across species, scales and experimental models, consistently distinguishing between conditions in which the brain state was systematically manipulated ^16,17^. Beyond this empirical robustness, PCI can be traced back to theoretical principles, and the neuronal underpinnings of its physiological and pathological alterations have been extensively studied ^16–22^. However, its relationship to broader principles of physics is unclear.

Like other living systems, the brain operates far from thermodynamic equilibrium ^23,24^. Neural dynamics are characterized by continuous energy dissipation, broken temporal symmetry, and directional information ^24–28^. Therefore, recent advances in non-equilibrium thermodynamics for whole-brain dynamics offer a promising mechanistic framework for addressing the conceptual limitations of complexity measures ^24,29^. In particular, the Fluctuation-Dissipation Theorem (FDT) states that in thermodynamic equilibrium, a system’s response to external perturbations is fully determined by the structure of its spontaneous fluctuations ^30^. In non-equilibrium systems, such as the brain, this balance no longer holds, and the degree to which FDT is violated can be used to quantify the system’s departure from equilibrium. Importantly, this violation is a marker of non-equilibrium dynamics and has been proposed and validated as a potential signature of consciousness in previous fMRI studies ^31,32^.

In this study, we hypothesized and empirically tested a theoretical link between PCI and FDT violations, proposing that both reflect a common underlying property of conscious brain states: their departure from thermodynamic equilibrium. Specifically, we suggest that the brain’s capacity to generate complex cortical responses to perturbations in conscious states is supported by a breakdown in the relationship between spontaneous fluctuations and evoked activity. From this perspective, high PCI values, which reflect rich and differentiated evoked responses, are predicted to coincide with strong FDT violations, indicating that the system output cannot be inferred from its intrinsic activity. Conversely, low PCI values observed in unconscious conditions are expected to reflect weaker violations of FDT, where the relationship between spontaneous and evoked activity is more preserved. Moreover, we hypothesized that FDT violations could be a promising complementary marker of consciousness, since they can be estimated from spontaneous activity alone, providing a mechanistically grounded, stimulation-free measure of consciousness.

To test these hypotheses, we analyzed neurophysiological recordings from animal models and human subjects across multiple levels of consciousness and experimental conditions. Specifically, evoked responses and spontaneous activity were recorded in mice under three different levels of anesthesia, as well as from two independent human EEG datasets. Among these two datasets, one included patients with disorders of consciousness (DoC), encompassing individuals diagnosed with unresponsive wakefulness syndrome (UWS) and minimally conscious states (MCS+, MCS−) with both vascular and traumatic etiologies. The second dataset served as a reference and included neurotypical individuals during wakefulness and under different anesthetic agents, enabling the generalization of our findings across conscious states and providing a brain-based comparison that does not rely on behavioral responsiveness. PCI was computed from cortical responses to direct perturbations, whereas FDT violations were assessed using a whole-brain model fitted to empirical recordings of spontaneous fluctuations.

The results show that FDT violations capture different levels of consciousness at a multiscale level and can discriminate among different states across all datasets. We also found a robust and systematic correlation in each species, dataset, and measurement modality: states with higher PCI values were associated with stronger FDT violations, whereas lower PCI values corresponded to smaller departures from equilibrium.

These findings provide further mechanistic foundation for PCI and other perturbation-based indices while introducing FDT violations as a complementary, perturbation-free measure for indexing consciousness. All in all, this study bridges systems and clinical neuroscience with the foundational principles of thermodynamic physics. This demonstrates that the complex dynamics associated with consciousness emerge from the intrinsic departure of the brain from equilibrium, reflecting thermodynamic asymmetries that characterize living systems. This framework offers a unifying perspective on consciousness grounded in non-equilibrium thermodynamics and opens new possibilities for assessing consciousness in basic research and clinical practice, as well as for understanding its neural substrates in both health and disease.

## RESULTS

Conscious brain states are thought to emerge from strong deviations from thermodynamic equilibrium, with a theoretical link to cortical complexity and PCI. To test this hypothesis, large-scale neurophysiological evoked and spontaneous recordings were analyzed from multiple species, modalities, and levels of consciousness. From these recordings, PCI is estimated from evoked cortical responses as a measure of complexity, whereas FDT violations are quantified from spontaneous activity to reflect the system’s non-equilibrium properties (Figure 1A).

**Figure 1.**
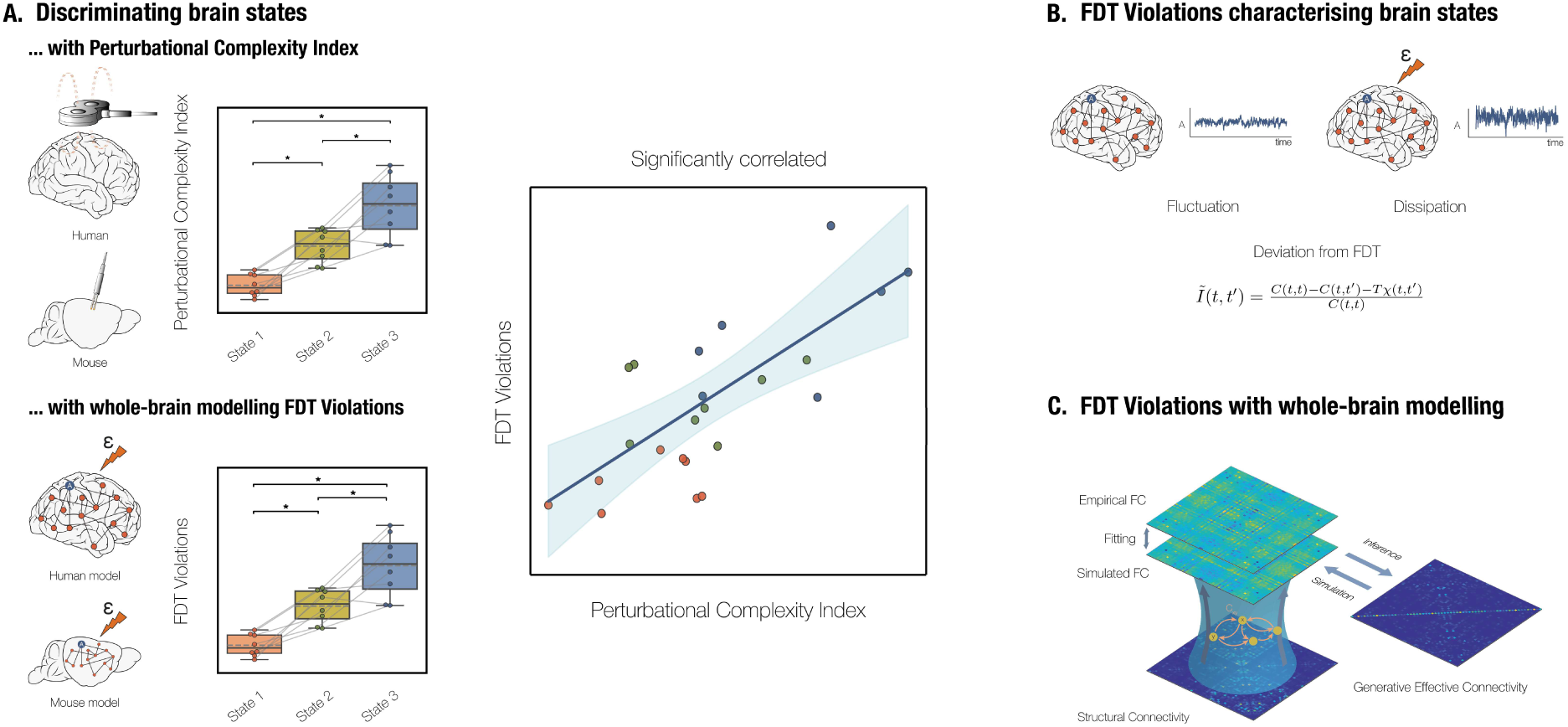
Framework for computing PCI and FDT violations from evoked and spontaneous brain dynamics. **(A)** PCI is computed from TMS-evoked or intracortical responses by binarizing spatiotemporal patterns of cortical source activations and quantifying their algorithmic complexity using Lempel-Ziv compression. PCI has proven to be a reliable and robust indicator, consistently distinguishing between conscious and unconscious conditions with high accuracy ^10,15^. FDT violations, estimated from spontaneous brain dynamics using whole-brain modelling, also discriminate between brain states and correlate strongly with PCI. This provides a mechanistic grounding for PCI and related perturbation-based indices, while also introducing FDT violation as a complementary, perturbation-free marker of consciousness. **(B)** Here, we use a model-based formalism for FDT violations that captures the mismatch between the model-derived response to a small theoretical perturbation and the response that would be expected if the system obeyed equilibrium. In equilibrium systems, fluctuations fully determine the response function; in the brain, departures from this relationship indicate non-equilibrium dynamics. FDT violations have been used in previous studies to characterize brain states ^31,32^. **(C)** Whole-brain modeling is applied at the individual level to reproduce spontaneous brain dynamics and capture the underlying asymmetrical structure of connectivity, referred to as Generative Effective Connectivity.

PCI values, derived from TMS-evoked or intracortical responses, were gathered from previous studies ^6,15,33,34^. In these datasets, spatiotemporal patterns of cortical source activations were binarized using non-parametric permutations, and the PCI was then computed using Lempel-Ziv compression algorithm. In parallel, we quantified FDT violations using a model-based formalism that captures the mismatch between the effective response to a small theoretical perturbation and the response that would be expected if the system obeyed equilibrium (Figure 1B). In equilibrium systems, fluctuations fully determine the response function; in the brain, departures from this relationship indicate non-equilibrium dynamics.

To obtain the effective response function, we used whole-brain models. First, the directed asymmetric interactions between brain regions were estimated by fitting a multivariate Ornstein-Uhlenbeck process to the empirical data. This approach has been successfully applied to characterize large-scale brain dynamics in both EEG and fMRI studies and has proven effective in extracting biologically meaningful structure from spontaneous activity ^35–37^. The model fitting was performed using a pseudo-gradient descent algorithm that simultaneously captures the zero-lag functional correlation matrix and the time-lagged covariance matrix, the latter encoding directional temporal interactions (Figure 1C). Once the model was fit, FDT violations were computed using the off-equilibrium extension provided in previous studies ^32,38,39^. After both PCI and FDT metrics were estimated, we assessed their correlation across conditions, as well as their relationship with clinical scores (Figure 1A).

We applied this pipeline across three distinct datasets: (i) invasive local field potentials (LFP) recordings from mice under graded anesthesia (N = 24), (ii) scalp EEG from neurotypical human participants in wakefulness and under three anesthetic agents (N = 30), and (iii) scalp EEG from patients with disorders of consciousness (N = 17). Within each dataset, we found that FDT violations increased from unconscious to conscious states and tightly co-varied with PCI, suggesting a unified mechanism underlying both spontaneous and evoked signatures of consciousness. These findings provide further mechanistic foundation for PCI and other perturbation-based indices while introducing FDT violations as a complementary, perturbation-free measure for indexing consciousness.

### FDT Violations Track Levels of Anesthesia in Animal Models

To investigate the relationship between PCI and FDT violations and to evaluate whether they can effectively track levels of consciousness, we first analyzed electrophysiological data from animal models. We examined large-scale LFP recordings from eight mice under three distinct levels of isoflurane anesthesia (N = 24). Both spontaneous and evoked brain activity were recorded.

As illustrated in Figure S1A, spontaneous activity during anesthesia was characterized by slow oscillations ^40^, reflecting the coordinated dynamics of large populations of neurons transitioning between active periods, known as Up states, and silent periods, referred to as Down states ^41,42^. The depth of anesthesia was associated with a slower dominant frequency and a higher occurrence of silent periods (See *Supplementary Note 1 & Supplementary Note 2)*.

PCI, derived from multi-unit activity–based evoked responses, significantly discriminated between all anesthesia conditions and consistently tracked changes in the anesthesia across levels (Spearman’s ρ = 0.71, *p* < 0.001; Figure 2A).

**Figure 2.**
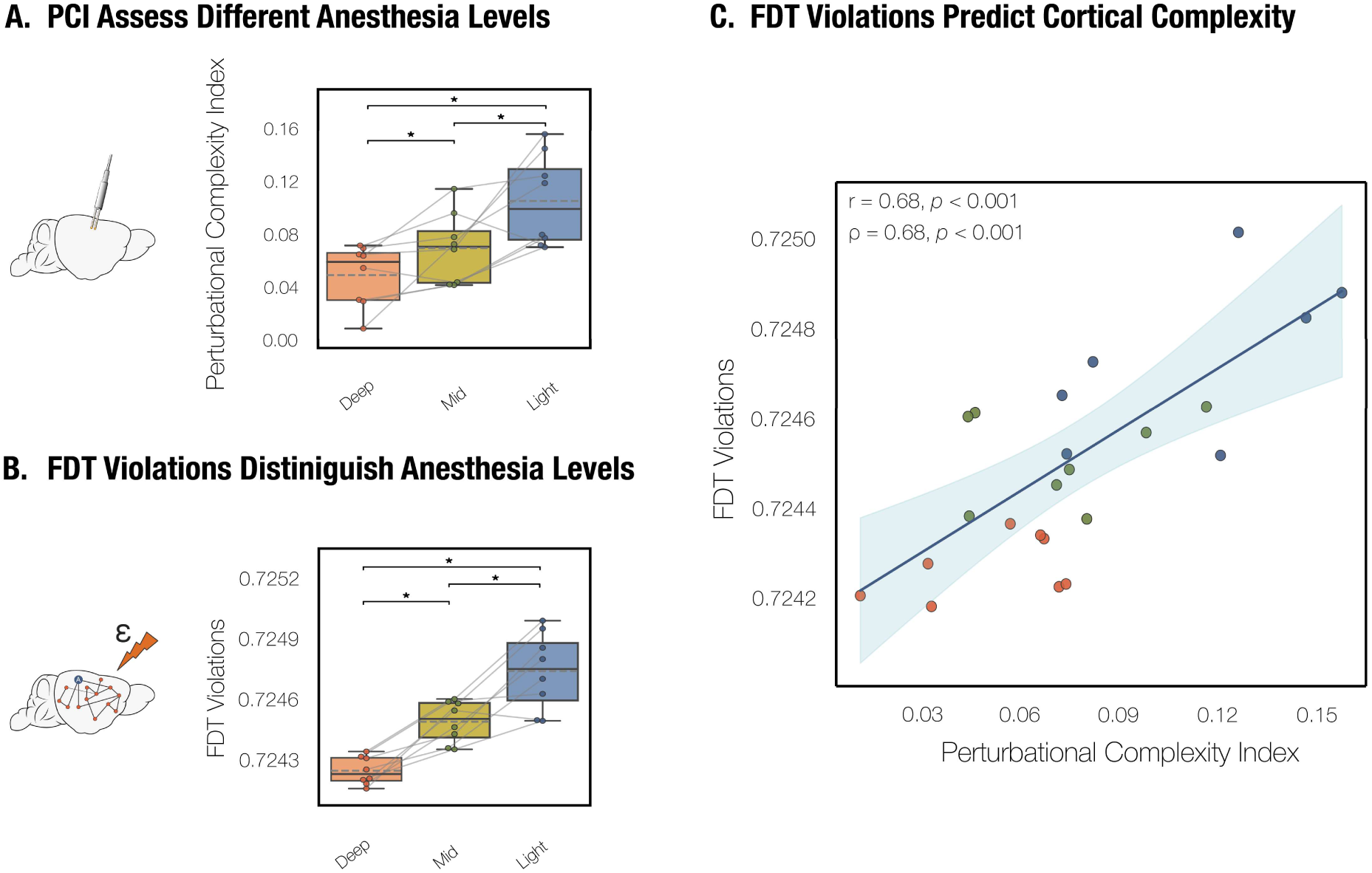
FDT violations track anesthesia levels in mice and correlate with the PCI. **(A)** PCI, calculated from multi-unit activity–based evoked responses, significantly discriminated between all anesthesia conditions. PCI consistently tracked the anesthesia levels in mice (Spearman’s ρ = 0.71, *p* < 0.001). **(B)** FDT violations calculated from the whole-brain model fitted to empirical spontaneous activity were also able to discriminate among all the conditions, as reflected by a strong monotonic relationship (Spearman’s ρ = 0.885, *p* < 0.001). **(C)** FDT violations correlate with the PCI (Pearson’s r = 0.68, *p* < 0.001; Spearman’s ρ = 0.68, *p* < 0.001). A linear mixed-effects model, controlling for anesthesia level and incorporating a random intercept for subject, confirmed that the association between remained significant (β = 0.613 ± 0.298 s.e.m., *p* = 0.037). All the pairwise comparisons were performed using Wilcoxon signed-rank tests and post hoc corrected with Holm-Bonferroni. * *p* < 0.05

In parallel, FDT violations, derived from whole-brain models fitted to the empirical spontaneous activity, also significantly distinguished all three levels of anesthesia (Figure 2B). This was supported by a robust monotonic relationship with anesthesia depth (Spearman’s ρ = 0.885, *p* < 0.001). FDT violations were also highly correlated with the dominant frequency of the slow oscillations (see *Supplementary Note 2*).

Figure 2C shows a strong correlation between PCI and FDT violations (Pearson’s r = 0.68, *p* < 0.001; Spearman’s ρ = 0.68, *p* < 0.001). A linear mixed-effects model, controlling anesthesia level and incorporating a random intercept for subject, confirmed that the association remained significant (β = 0.613 ± 0.298 s.e.m., *p* = 0.037).

### FDT Violations Distinguish Wakefulness from Anesthesia in Human EEG

To generalize the findings to humans, we analyzed EEG recordings from neurotypical individuals. These data enabled us to extend the results across distinct conscious states and evaluate our findings with brain-based markers of consciousness that do not rely on behavioral responsiveness. The dataset included fifteen healthy participants recorded during wakefulness, with eyes-open or eyes-closed conditions. Participants were randomly assigned to one of three anesthetic conditions (N = 5 each for xenon, propofol, and ketamine), resulting in a total of N = 30 recordings. Participants under ketamine anesthesia reported vivid, dream-like experiences during unresponsiveness, consistent with previous studies describing ketamine-induced conscious states ^6,43,44^.

As illustrated in Figure S1B, spontaneous activity during anesthesia was dominated by slow oscillations, while wakefulness was characterized by prominent alpha rhythms ^22,45^.

PCI successfully distinguished participants who retrospectively reported any form of conscious experience from those who did not. To evaluate its performance as a binary classifier, the area under the receiver operating characteristic (AROC) curve was computed, finding that PCI achieved an AROC of 1.0, indicating perfect classification between conscious and unconscious states in this sample. A strong monotonic relationship was also observed between PCI and conditions (Spearman’s ρ = 0.867; *p* < 0.001), indicating that PCI could index consciousness during anesthesia conditions (Figure 3A). PCI did not differentiate the ketamine condition from wakefulness, which aligns with the presence of ongoing conscious activity despite behavioral unresponsiveness.

**Figure 3.**
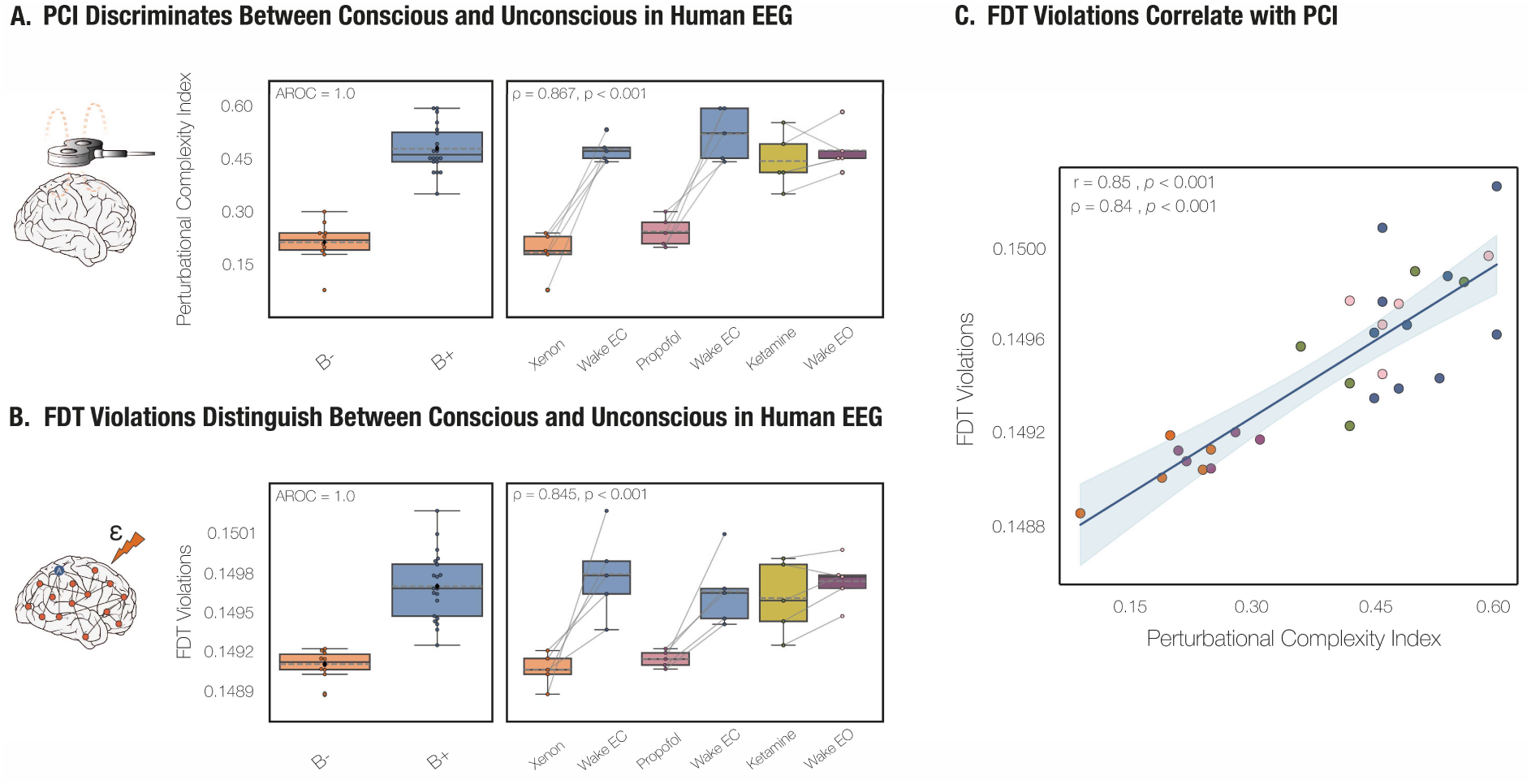
FDT violations in human EEG track level of consciousness during wakefulness and anesthesia and correlate with PCI. **(A)** PCI discriminated between participants who retrospectively reported some form of conscious experience (B+) and those who did not (B-). PCI achieved a perfect classification performance (AROC = 1.0). A strong monotonic relationship with condition was observed (Spearman’s ρ = 0.867, *p* < 0.001). **(B)** FDT violations, computed from whole-brain models fitted to spontaneous EEG activity, also differentiated participants who reported conscious experience from those who did not (AROC = 1.0). A monotonic relationship was observed (Spearman’s ρ = 0.845, *p* < 0.001), consistent with sensitivity to differences in conscious state. **(C)** FDT violations were strongly correlated with PCI (Pearson’s r = 0.85, *p* < 0.001; Spearman’s ρ = 0.84, *p* < 0.001). A linear mixed-effects model, controlling for anesthesia state and drug type while including subject identity as a random intercept, confirmed that the association between FDT violations and PCI remained significant (β = 0.564 ± 0.275 s.e.m., *p* < 0.001).

FDT violations also distinguished participants who reported conscious experience from those who did not. This was supported by an AROC of 1.0 and a similarly strong monotonic correlation (Spearman’s ρ = 0.845; *p* < 0.001; Figure 3B). As with PCI, FDT violations did not separate the ketamine condition from wakefulness.

FDT violations were strongly correlated with PCI (Pearson’s r = 0.85, *p* < 0.001; Spearman’s ρ = 0.84, *p* < 0.001; Figure 3C). A linear mixed-effects model, controlling for anesthesia state (anesthetized vs. awake) and drug type, while including subject identity as a random intercept, confirmed that the association between FDT violations and PCI remained significant (β = 0.564 ± 0.275 s.e.m., *p* < 0.001).

### FDT Violations Correlate with PCI and Clinical Scores in Disorders of Consciousness

Lastly, to evaluate the clinical utility of our findings in the context of disorders of consciousness, we analyzed EEG data from patients diagnosed with chronic DoC, including individuals classified as UWS, MCS−, and MCS+. Clinical diagnosis was based on repeated CRS-R assessments conducted at least three times within one week. EEG recordings were obtained from 17 patients, with the cohort comprising individuals with vascular (N = 13) and traumatic (N = 4) etiologies.

Consistent with previous reports, PCI reliably distinguished between behaviorally responsive and unresponsive patients (Cliff’s *δ* = 1.0, *p* < 0.001; Figure 4A). To evaluate its performance as a binary classifier, the AROC was computed. PCI reached a value of 1.0, indicating perfect classification between conscious and unconscious states in this sample.

**Figure 4.**
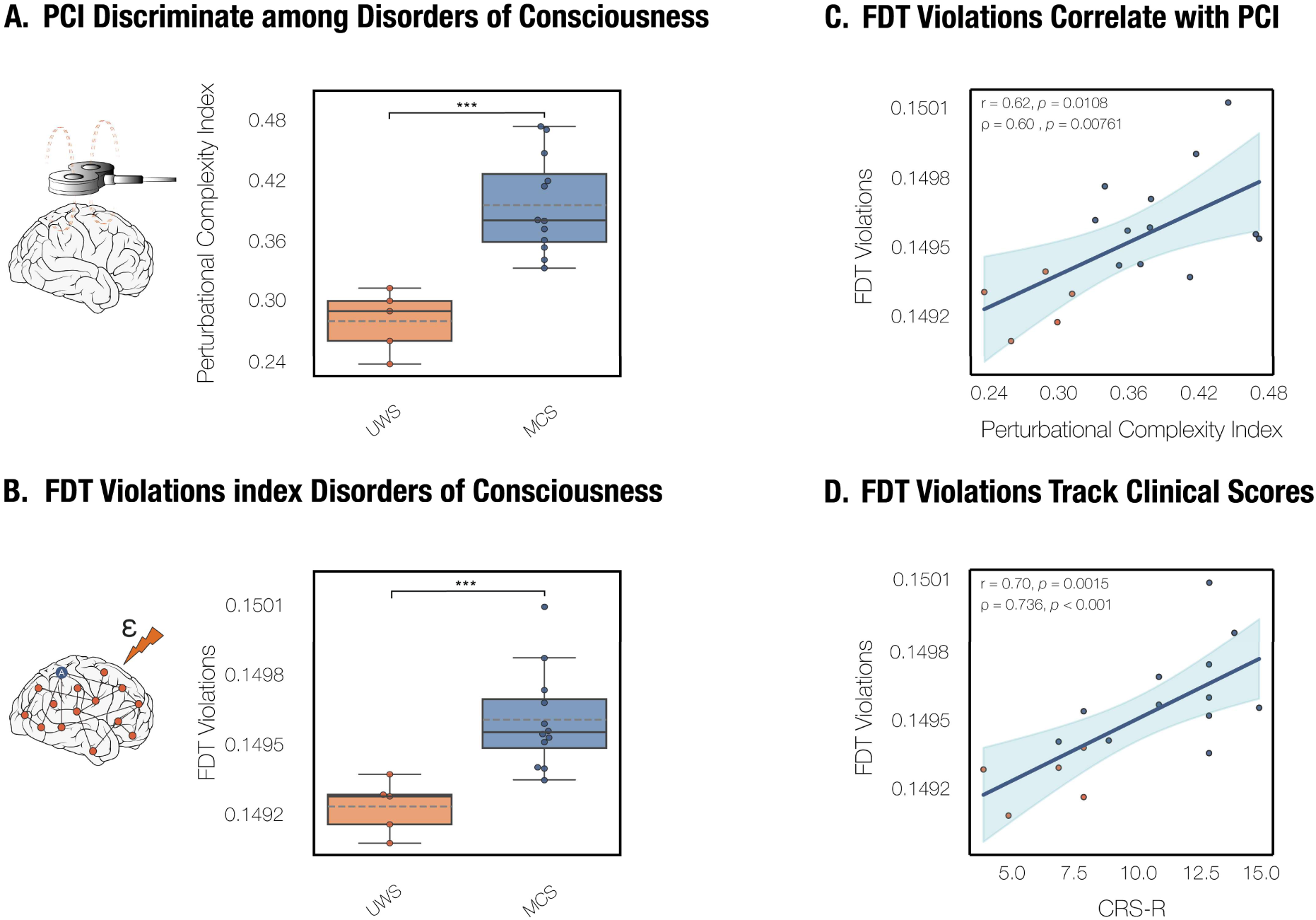
FDT violations distinguish conscious from unconscious states in patients with disorders of consciousness and correlate with PCI and clinical scores. **(A)** PCI discriminated between conscious (MCS-/MCS+) and unconscious patients (UWS) with high accuracy (Cliff’s *δ* = 1.0, *p* < 0.001) and achieved perfect classification performance (AROC = 1.0). **(B)** FDT violations also distinguished conscious (MCS-/MCS+) from unconscious (UWS) patients with similarly high accuracy (Cliff’s *δ* = 0.967, *p* < 0.001) and achieved an AROC of 0.967. **(C)** FDT violations were significantly correlated with PCI values, indicating that both metrics capture related aspects of conscious in a clinical context (Pearson’s r = 0.62, *p* = 0.0108; Spearman’s ρ = 0.60, *p* = 0.0076). An ANCOVA including the *FDT violations × Condition* interaction term revealed no evidence of slope heterogeneity between groups, indicating that the relationship was conserved across conditions (F(1,13) = 0.007, *p* = 0.934; Freedman–Lane permutation *p_Perm_* = 0.908). **(D)** FDT violations were also strongly correlated to clinical scales such as CRS-R scores (Pearson’s r = 0.707, *p* = 0.0015; Spearman’s ρ = 0.736, *p* < 0.001). All p-values were calculated using two-sided Mann-Whitney U test as the groups were independent. *** *p* < 0.001.

FDT violations also significantly differentiated behaviorally responsive from unresponsive patients (Cliff’s *δ* = 0.967, *p* < 0.001; Figure 4B), offering a complementary robust marker of consciousness that does not require external perturbation. The corresponding AROC was 0.967, confirming excellent discriminative performance in this clinical setting.

In this more challenging clinical context, FDT violations remained significantly correlated with PCI (Pearson’s r = 0.62, *p* = 0.0076; Spearman’s ρ = 0.60, *p* = 0.0108; Figure 4C). An ANCOVA including *FDT violations × Condition* interaction term revealed no evidence of slope heterogeneity between groups, indicating that the relationship was conserved across conditions (*F*(1,13) = 0.007, *p* = 0.934; Freedman–Lane permutation *p_perm_* = 0.908). Notably, FDT violations correlated more strongly with the CRS-R scores (Pearson’s r = 0.707, *p* = 0.0015; Spearman’s ρ = 0.736, *p* < 0.001; Figure 4D), than PCI did (Pearson’s r = 0.56, *p* = 0.018; Spearman’s ρ = 0.57, *p* = 0.0116).

## DISCUSSION

Here, we show that FDT violations provide a link between PCI and fundamental principles of physics, potentially paving the way to the implementation of novel model-based markers of consciousness.

Across three different electrophysiological datasets in mice and humans, FDT violations captured differences in consciousness levels and reliably distinguished between multiple conscious states, including wakefulness, general anesthesia, ketamine-induced unresponsiveness, and DoC. Moreover, we found a robust and systematic correlation between FDT violations and PCI in all datasets, species, and modalities. Together, these results propose and establish a unified framework linking spontaneous dynamics, perturbational complexity, and fundamental thermodynamic principles.

Currently, the assessment of consciousness relies on behavioral responsiveness, most commonly assessed with the CRS-R scale in the clinical context. However, an absence of responsiveness cannot be considered as absence of consciousness ^3–5^. Motor, sensory, or executive impairments may prevent behavioral interaction with the environment, even when awareness is preserved, as in healthy subjects during dreaming, under general anesthesia, or in cases of severe brain injury ^7–9,46^. This limitation has motivated the search for non-behavioral markers of consciousness. Early candidates such as the P3b component, global EEG activation, and cortical oscillations showed initial promise but have proven unreliable ^47–49^. In practice, they failed to consistently separate minimally conscious from unresponsive patients or to index consciousness properly across altered states ^33,50–54^.

The development of new quantitative complexity indices marked an important advance, inspired by theoretical neuroscience and particularly by Integrated Information Theory ^13,55–57^. Empirically, brain complexity has been quantified either through analysis of spontaneous activity ^58–61^ or by quantifying reactivity patterns to cortical perturbations ^6,10^. Among these, PCI has emerged as a robust and generalizable marker, reliably distinguishing conscious from unconscious conditions (As also seen here in Figure 2A, Figure 3A & Figure 4A) ^10,11,15^. While PCI can be traced back to theoretical principles ^16–22^, its relationship to broader principles of physics remains unclear.

Our study directly addresses this open question by providing a novel empirical link between PCI and FDT violations across species, brain states and modalities (Figure 2C, Figure 3C, Figure 4C). Specifically, we find that states with higher PCI values, reflecting richer, more differentiated evoked responses, are associated with stronger FDT violations, while states with lower PCI values correspond to more equilibrium-like dynamics. These findings offer empirical support for a theoretical connection between brain complexity and non-equilibrium thermodynamics, giving a proper explanation to the successful findings from previous studies through a non-equilibrium perspective ^6,10,15,16,34^. This interpretation is further supported by complementary analyses of other non-equilibrium measures, including entropy production, temporal irreversibility, and coupling asymmetry, which showed parallel reductions with loss of consciousness (see *Supplementary Note 3*). In this view, both PCI and FDT violations may reflect a common underlying property: the system’s departure from thermodynamic equilibrium. Mechanistically, the brain’s capacity to generate complex, integrated cortical responses in conscious states is supported by a breakdown in the relationship between spontaneous fluctuations and evoked activity. This breakdown, captured as FDT violations, indicates that the system’s output cannot be fully predicted from its intrinsic fluctuations alone. In contrast, unconscious states, characterized by lower PCI, exhibit weaker FDT violations, suggesting a more constrained, predictable mapping between spontaneous and evoked dynamics.

Previous studies have reported relationships between PCI and emergent physics phenomena such as criticality ^62^, as well as modeling links between PCI and irreversibility arising from non-equilibrium dynamics ^63^. Here, we provide the first empirical evidence that complexity-based evoked measures are related to direct estimates of FDT violations, a fundamental principle of physics governing equilibrium and its breakdown. This correspondence links conscious dynamics to non-equilibrium fluctuations, strengthening the theoretical foundation of complexity-based approaches and offering a complementary thermodynamic perspective on current theories of consciousness.

Our study also demonstrated that FDT violations could constitute a promising, complementary marker of consciousness, applicable to electrophysiological recordings in both humans and animal models. FDT violations successfully distinguished three levels of anesthesia in mice LFP data (Figure 2B). In humans, FDT violations reliably indexed states of wakefulness and anesthesia induced by xenon and propofol (Figure 3B). However, similar to PCI, they did not distinguish between wakefulness and the dissociative state induced by ketamine at anesthetic concentrations ^6^. This observation aligns with prior clinical studies and with participants reporting “ketamine dreams”, rich inner experiences upon awakening, often unrelated to their surgical environment ^6,43,44^. These results suggest that ketamine-induced conscious states preserve cortical complexity and remain far from thermodynamic equilibrium. In patients with DoC, FDT violations were able to significantly differentiate between unresponsive (UWS) and minimally conscious (MCS+/MCS−) patients (Figure 4B) and correlated with CRS-R scores (Figure 4D). Overall, FDT violations captured meaningful distinctions in brain states across all datasets and at multiple scales, introducing a complementary, model-based approach to index consciousness.

Nonetheless, future work should assess the generalization of this methodology to more complex clinical populations, such as severe post-anoxic patients, who are readily identified by TMS-EEG as showing no significant response ^15^. In such cases, refined modeling may be required to infer asymmetrical connections and reliably estimate FDT violations, as these individuals often present distinct neurophysiological signatures, including globally reduced cortical activity leading to low-voltage EEG patterns ^64–67^. This framework can also be used to capture focal alterations in brain dynamics. Following brain lesions, for example, quantifying local FDT violations could yield spatially resolved maps of non-equilibrium dynamics, offering mechanistic insight into how injury reshapes integration and pointing to potential targets for restoring wake-like activity through neuromodulation ^68^. Taken together, this perspective suggests that FDT violations may offer a powerful, non-invasive, and generalizable approach for mechanistic insight into a wide range of neurological conditions.

Moreover, importantly, the relationship between FDT violations and fundamental neural dynamics remains an open area of investigation. In fact, the emergence of spontaneous slow oscillations (SO), a hallmark of low-complexity brain states, is commonly observed during non-rapid eye movement sleep, in patients with UWS, even during periods of eyes open, and in other pathological states ^45,53,58,69^. These oscillations reflect cortical bistability as they are characterized by alternating phases of neuronal tonic firing and neuronal silence ^70^. Experimental and computational studies across humans, rodents, and cortical slice preparations converge in identifying cortical bistability and OFF-periods as key disruptors of large-scale network integration and complexity ^16–19,22^. Cortical perturbations reveal that bistability causally constrains network dynamics ^17–19,22^. PCI reflects this directly, increasing with rich, specific interactions and decreasing when OFF-periods yield restricted, stereotypical responses^10^. Theoretically, FDT violations provide a natural framework to study this phenomenon. They quantify the breakdown in the relationship between spontaneous fluctuations and evoked activity, thereby capturing the brain’s capacity to generate complex, integrated responses—an essential feature of conscious states It should be noted that this study has several limitations. First, electrophysiological measures can be affected by artifacts or limited electrode coverage, although standard preprocessing minimizes such effects. In DoC patients, diagnostic reliability is further challenged by behavioral fluctuations and heterogeneous signal quality; while the CRS-R vigilance protocol mitigated this, future work should examine robustness in larger and more diverse cohorts ^71,72^. Time since injury may also influence EEG features, but prior studies suggest only marginal effects, and our focus was on diagnostic capacity rather than longitudinal progression ^73^. Methodologically, we employed linear stochastic dynamics for analytical tractability, which may underestimate nonlinear aspects of brain activity. FDT violation values should be interpreted cautiously across species or recording modalities, given differences in spatial resolution, signal properties, and recording conditions. The limited number of subjects per condition also constrains the precision of correlation and mixed-model estimates. Despite these caveats, the consistency of results across species, brain states, and modalities underscores the robustness of the main findings, but further validation in larger and more varied populations will be essential. It is also worth noting that while here we implemented a model-based whole-brain framework, similar approaches could be directly applied to empirical brain dynamics under linear stochastic assumptions ^74^. Importantly, the model-based perspective remains advantageous as it allows for in silico manipulations, offering a path to bridge empirical data with mechanistic theory.

All in all, this study identifies FDT violations as a complementary, reliable, mechanistically grounded, and stimulation-free marker of consciousness, consistent across species, recording modalities, and scales. Most importantly, by establishing a systematic relationship between FDT violations and PCI across brain states and conditions, we demonstrate that perturbational complexity and non-equilibrium dynamics are closely related. Finally, this study bridges brain complexity and non-equilibrium dynamics, grounding consciousness in fundamental principles of physics. Together, these findings introduce a unifying framework in which spontaneous brain dynamics, perturbational complexity, and non-equilibrium thermodynamics converge to explain brain states. This perspective opens new avenues for understanding the nature of consciousness and for developing objective tools to assess its loss and recovery in health and disease. These findings also lay the conceptual foundation for future efforts to identify strategies capable of restoring consciousness.

## METHODS

### Mice Electrophysiological Data

#### Animal Subjects

Data were obtained from a previously published study ^34^. Recordings were conducted in eight adult male C57BL/6J mice bred in-house at the University of Barcelona and maintained on a 12 h light/dark cycle with ad libitum access to food and water. All procedures were approved by the Ethics Committee of the Hospital Clinic of Barcelona and adhered to Spanish regulatory laws (BOE 34/11370-421, 2013) and the European Union directive 2010/63/EU. Detailed methodological procedures can be found in the original publication ^34^.

#### Surgical Procedures and Anesthesia Protocol

Anesthesia was induced via intraperitoneal injection of ketamine (75 mg/kg) and medetomidine (1.3 mg/kg) and maintained with isoflurane in pure oxygen. Atropine (0.3 mg/kg), methylprednisolone (30 mg/kg), and mannitol (0.5 g/kg) were administered subcutaneously to reduce respiratory secretions and prevent cerebral edema. Body temperature was monitored and maintained at 37°C using a thermal blanket (RWD Life Science, China).

Mice were placed in a stereotaxic frame (SR-6M, Narishige, Japan), and a craniotomy and durotomy (−3.0 mm to +3.0 mm relative to bregma and +3.0 mm lateral to midline) was performed over one hemisphere. Three anesthesia levels were defined based on isoflurane concentrations: Deep = 1.16% ± 0.08 s.e.m.; Medium = 0.34% ± 0.06 s.e.m.; and Light = 0.1–0%. The volume delivered was 0.8 L/min and the animals were breathing freely via a tracheostomy. Each level was maintained for 20–30 min, and recordings were taken during a stable slow-oscillatory regime (∼10 min post-concentration change, before microarousals) ^75^. Reflexes were regularly tested to confirm anesthetic depth.

#### Electrophysiological Recordings

Extracellular LFP activity was recorded using 32-channel multielectrode arrays (550 μm interelectrode spacing) covering the entire exposed area of the cortex ^76^. Signals were amplified by 100, high-pass filtered above 5 kHz, digitized at 5 kHz, and fed into a computer via a digitizer interface (CED 1401 and Spike2 software, Cambridge Electronic Design, UK).

For each anesthesia level, 300–500 seconds of spontaneous activity was recorded, followed by an equal duration of evoked activity used for PCI calculation. For further methodological details on the stimulation setup and protocol, readers are referred to the original publication where these procedures are described in detail ^34^.

### Human EEG Data During Wakefulness and Anesthesia

#### Participants

Data were obtained from a previously published dataset ^77^ comprising fifteen healthy participants (5 males, aged 18–28 years) recorded during wakefulness with eyes-open or eyes-closed conditions. Participants were randomly assigned to one of three anesthetic conditions (N = 5 per condition: xenon, propofol, and ketamine), resulting in a total of N = 30 recordings ^73^. All participants underwent medical screening to rule out contraindications to anesthesia and provided written informed consent. The experimental protocol was approved by the Ethics Committee of the University of Liège, Belgium. This dataset has been validated and widely used in prior research ^15,62,73^. For full methodological details, refer to the original publication ^73^.

#### Experimental Design and Anesthetic Procedures

All experimental procedures were performed at the Centre Hospitalier Universitaire in Liège, Belgium. Spontaneous EEG, followed by TMS-EEG recordings, was recorded before drug administration (Ramsay Scale score 2) as well as upon reaching deep unresponsiveness in three consecutive assessments (Ramsay Scale score 6) ^78^. At the end of the TMS-EEG recordings, anesthesia was discontinued, participants were allowed to recover, and assessments of responsiveness were continued every 30 seconds. Participants were continuously monitored for heart rate, oxygen saturation, blood pressure, exhaled CO₂, and temperature. Mild nausea, if present, was treated with metoclopramide (2 mg).

To enable comparisons across drugs with distinct molecular targets, all anesthetic protocols aimed to reach a common behavioral endpoint: unresponsiveness to external stimuli systematically assessed using repeated Ramsey Scale administrations. Anesthetic protocols followed established procedures for propofol ^79^, xenon ^80^, and ketamine ^81^.

#### TMS–EEG Acquisition

EEG data were recorded using a 60-channel TMS-compatible amplifier (Nexstim Plc., Finland), with online filtering (0.1–350 Hz) and sampling at 1450 Hz. The reference electrode was placed on the forehead, and the impedance was kept below 5 kΩ. Two additional channels recorded electrooculographic activity. For further methodological details on the TMS–EEG acquisition, stimulation, and navigation procedures, readers are referred to the original studies where these methods are described in detail ^73^.

#### Assessment of Conscious Experience

To assess the presence or absence of conscious experience during anesthesia-induced unresponsiveness, retrospective reports were collected in all participants after awakening. Responsiveness was considered restored when participants consistently reached a Ramsay score of 2 on three consecutive assessments. At that point, they were asked whether they recalled any mental activity during the unresponsive state. Conscious experience was defined broadly as any kind of mental activity, including thoughts, dreams, images, or emotions. Under propofol and xenon, participants either reported no memory of the period or were unable to recall any experience. In contrast, all participants who received ketamine described vivid, emotionally intense experiences with a coherent narrative structure, consistent with previous findings ^6^.

### Human EEG Data in Disorders of Consciousness

#### Patients

Data were gathered from previously published studies ^15,33^ and included a subset of 17 patients diagnosed with prolonged or chronic DoC. Patients were clinically classified as being in UWS, MCS−, or MCS+ based on repeated CRS-R assessments conducted at least three times within one week. The cohort comprised patients with vascular (N = 13) and traumatic (N = 4) etiologies. Patients with post-anoxic etiology were not included in this study because, in severe cases, their electrophysiological signatures primarily reflect cortical integrity rather than consciousness capacity^33^. These patients often show marked suppression of alpha power within a context of overall power suppression and low-voltage EEG ^64–67,82,83^.

Written informed consent was obtained either from the patients (when possible) or from their legal representatives. All experimental procedures were approved by the respective local ethics committees. All patients were free of sedative medication at the time of recording. Further methodological details can be found in the original publications ^15,33^.

#### Experimental Design

During recordings, patients were seated in an upright or reclined position in a quiet room. Short spontaneous EEG recordings (median duration = 5.13 minutes, interquartile range: 4.03–6.59 minutes) were acquired either during the same experimental session of TMS-EEG recordings (N = 14) or, when not feasible, during a closely timed session within one day (N = 1) or at most within two weeks (N = 2). In all cases, the CRS-R behavioral diagnosis remained stable across sessions. Patients were continuously monitored by trained personnel to detect any signs of drowsiness. When necessary, the CRS-R vigilance protocol ^2^ was administered to minimize the risk of sleep intrusions and reduce the potential confounding effects of arousal fluctuations on the variables of interest.

In the DoC population, behavioral classification based on CRS-R yielded a natural dichotomy: behaviorally unresponsive patients (B−, i.e., UWS; N = 5) and behaviorally responsive patients (B+, i.e., MCS− and MCS+; N = 12).

#### TMS-EEG Acquisition

EEG was recorded using Ag/AgCl electrodes and TMS-compatible amplifiers (Nexstim Plc., 60 channels, N = 14; Brain Products GmbH, 64 channels, N = 3). The two systems employed highly similar electrode montages (see *Supplementary Note 4*), with a common acquisition reference placed on the forehead near Fpz. Electrode input impedance was maintained below 20 kΩ. For further methodological details on the TMS–EEG acquisition, stimulation, and navigation procedures, readers are referred to the original studies in which these methods are described in detail ^15,33^.

### Preprocessing

#### Animal Electrophysiological Signals

For rodent data, distinct signal components were extracted from the same LFP recordings. For PCI computation, multiunit activity was estimated by computing the power in the 200–1500 Hz band using 5-ms windows ^40,84,85^. This spectral range captures local neuronal firing and primarily reflects fast, evoked population spiking, making it well suited for quantifying spatiotemporal complexity without contamination ^86^. Further methodological details can be found in the original publication ^34^.

In contrast, FDT violations analysis required signals that reflect the brain’s spontaneous, large-scale dynamical regime. To this end, LFPs were bandpass filtered between 0.1 and 4 Hz to isolate slow cortical oscillations, which dominate anesthesia (accounting for up to 95% of spectral power) and are particularly sensitive to global state changes. ^41,70^. These slow fluctuations evolve over extended timescales and display rich temporal and spatial structure, providing a natural substrate for assessing FDT violations, which quantify a system’s departure from equilibrium. Importantly, this band choice was further supported by repeating the analysis using the broadband range employed for human data (see *Supplementary Note 5*).

#### Human EEG signals

Unlike in rodents, human recordings spanned a broader range of brain states, including wakefulness, that involve additional spectral components such as alpha and beta rhythms. For this reason, broadband signals were preserved to capture the full repertoire of state-dependent dynamics.

Spontaneous EEG signals were bandpass filtered using a third-order, zero-phase Butterworth filter (0.5–60 Hz cutoffs) with the *filtfilt* function in MATLAB, and notch-filtered to remove 50 Hz line noise and its harmonics. Data were then downsampled to reduce computational demands (from 5000 Hz to 1000 Hz for Brain Products recordings, and from 1450 Hz to 725 Hz for Nexstim recordings).

A trained neurophysiologist manually inspected each recording to exclude segments contaminated with gross artifacts. This procedure retained a median of 4.79 minutes of clean data per condition (98.74%, interquartile range: 3.58–6.12 min, 91.06–99.27%) and excluded a median of 2 electrodes per subject (IQR: 1–4). Rejected electrodes were interpolated using spherical splines, and all signals were re-referenced to the common average. To remove residual non-neural activity, independent component analysis was performed. Components attributable to ocular, muscular, or cardiac artifacts were visually identified and excluded. A median of 33 components (IQR: 25–41.5) were retained and back-projected to reconstruct artifact-free EEG signals. Further details on recording setups and artifact removal procedures are provided in *Supplementary Note 4*.

For FDT computation, the cleaned EEG data were low-pass filtered at 40 Hz and downsampled to the corresponding Nyquist rate to minimize high-frequency muscle artifacts and reduce computational demands during whole-brain model fitting. These preprocessing steps ensured high signal fidelity and minimized non-neural confounds, preserving the information necessary for for the calculation of FDT violations.

For methodological details on the TMS-EEG preprocessing, please refer to the original studies ^6,15,33^. For illustration, representative TMS–EEG traces corresponding to high and low PCI values are shown in *Supplementary Note 6*.

### Perturbational Complexity Index

PCI values were obtained from previous works ^6,15,33,34^. PCI quantifies the complexity of spatiotemporal brain patterns in response to external perturbations, typically delivered via TMS or intracranial stimulation. Complexity is measured using the lossless Lempel-Ziv (LZ) data compression algorithm, normalized by the source entropy ^10^. To compute the LZ complexity, TMS-evoked responses are first binarized, producing a significant spatiotemporal matrix. This is achieved using a non-parametric permutation statistical procedure, as described by Casali et al. (2013) ^10^. The maximum PCI value across all stimulation sites (PCImax) is typically retained as the final score. Further methodological details for PCI computation in both humans and animal models can be found in the original studies ^6,15,33,34^.

It should be noted that when PCI is applied to intracranial recordings and multiunit activity, particularly in animal models, it is often referred to as intracranial multiunit PCI (imPCI). This variant extends the original index to accommodate multiunit spike-based and intracortical perturbation paradigms ^16,34^.

### Whole-Brain Modelling

The local dynamics of each brain region were modeled as a multivariate Ornstein–Uhlenbeck (OU) process, which corresponds to the linearization of nonlinear population models such as Wilson–Cowan neurons ^88^. This formulation provides a linear approximation to the stochastic dynamics of the system near a stable fixed point ^89,90^. OU processes are particularly well-suited for capturing non-equilibrium dynamics, as they enable a direct and quantitative assessment of the relationship between spontaneous fluctuations and linear response functions, allowing deviations from equilibrium to be computed ^35,38,91^. Moreover, OU-based models have been successfully applied to characterize large-scale brain dynamics in both EEG and fMRI studies, and have proven effective in extracting biologically meaningful structure from spontaneous activity ^35–37^.

In this framework, brain activity is modeled as a network of interacting regions governed by coupled stochastic differential equations:

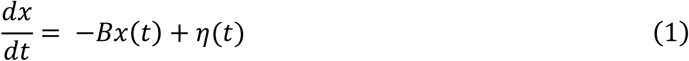

where *B* is the drift matrix and corresponds to the asymmetric connectivity matrix inferred from empirical data. The *η*(*t*) term corresponds to zero-mean additive white Gaussian noise. The noise is characterized by its covariance structure, < *η*(*t*)*η*(*t*’) > = 2 *σ*^2^*δ*(*t* − *t*’), where *σ*^2^ is, by definition, symmetric and positive semidefinite.

To ensure a well-defined and stable stationary solution, we require that all eigenvalues of the drift matrix *B* have strictly positive real parts ^90^. This condition guarantees that the associated system dynamics fulfils the Hurwitz criterion, meaning the system naturally decays and converges to a unique steady state. This requirement is essential not only for mathematical stability and well-posedness, but also for the thermodynamic and biological interpretability of the model ^90^. The model describes multivariate neural activity signals while preserving the temporal structure and directional interactions required to quantify non-equilibrium dynamics and assess how departures from equilibrium relate to changes in consciousness. This approach is consistent with the methodology and previous applications of the model ^36,92–94^.

### Model Optimization

To quantify FDT violations, we applied the multivariate OU modeling framework described in the section above. The empirical observables used for model fitting included the functional connectivity matrix (*FC^empirical^*), reflecting zero-lag statistical dependencies across brain regions, and the normalized time-lagged covariance matrix (*FS^empirical^*(*τ*)), which captures directional temporal interactions at a fixed lag while reducing zero-lag dependencies and volume-conduction effects. These normalized time-lagged covariance matrices are generated by taking the shifted covariance matrix *KS^empirical^*(*τ*) and normalizing each pair (*i*, *j*) by dividing 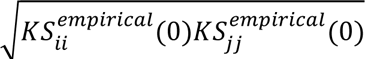. This normalization accounts for variance across regions and emphasizes relative lagged dependencies. Importantly, incorporating time-lagged covariances allows the estimated coupling matrix *B* to become asymmetric, enabling the emergence of non-equilibrium dynamics and quantifiable violations of the FDT ^95,96^. The time lag *τ* was selected to ensure theoretical consistency with the OU formulation and to preserve sensitivity to temporal asymmetries, following previous studies ^35,37^.

To match these empirical observables, the model was iteratively adjusted using a heuristic pseudo-gradient descent algorithm that minimized the difference between simulated and empirical *FC* and *FS*(*τ*) matrices ^37,97^. We updated *B* until the fit was fully optimized and converged to a stable solution. More specifically, the updating uses the following rule:

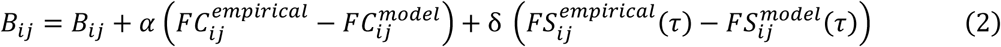

The multivariate OU model allows analytic computation of both stationary and time-lagged covariances. The stationary covariance matrix was defined as *K* = 〈*xx^T^*〉 and the noise covariance as *Q*, assumed to be symmetric and positive semidefinite. Using Itô’s calculus and keeping terms to first order in dt, the evolution of *K* is given by the following equation ^31,32,96^:

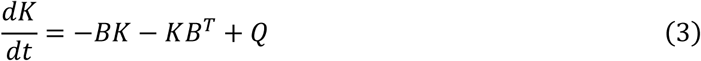

At the steady state 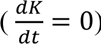, we obtain the continuous-time Lyapunov equation:

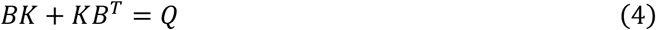

Solving this equation (e.g., via Lyapunov or Sylvester solvers) provides the model-derived covariance matrix *K*, from which the simulated functional connectivity (*FC^model^*), is computed. The time-lagged covariance was then calculated analytically as:

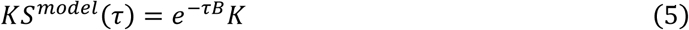

and normalized in the same manner as the empirical data. Note that *K*(0) = *K*.

*B* initialized as a random symmetric matrix constrained to satisfy the Hurwitz stability criterion. This approach ensured mathematical and thermodynamic stability without imposing anatomical priors. Learning rates were fixed at *α* = *δ* = 0.00001, and the optimization was repeated until the solution stabilized. The full optimization process was repeated 1000 times, and the results were then averaged to ensure convergence to a robust solution. We refer to the final optimized matrix as the generative effective connectivity matrix, which captures both the directional and non-equilibrium aspects of spontaneous brain dynamics ^98^. Noise was assumed to be homogeneous and therefore did not need to be fitted, as demonstrated in *Supplementary Note 7*.

### FDT Violations

To assess the departure from equilibrium, we adapted a model-based formalism grounded in the off-equilibrium extension of FDT ^38,39,99^. In thermodynamic equilibrium, the FDT states that the system’s linear response to small perturbations is entirely determined by spontaneous fluctuations. However, in systems that are out of equilibrium, such as the brain, this relationship is broken. The degree of FDT violation therefore provides a principled and physically interpretable measure of non-equilibrium dynamics ^31,32^.

More precisely, in equilibrium, the linear response function *R*(*t*, *t*’) of the system’s state *x*(*t*) to a small enough perturbation *∊*(*t*’) is defined as ^30^:

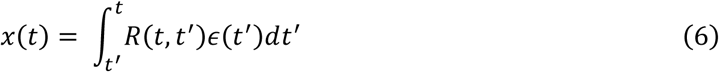

According to the FDT, this response function is directly related to the autocorrelation function of spontaneous fluctuations, *C*(*t*, *t*’) = 〈*x*(*t*)*x*(*t*’)〉, through the expression:

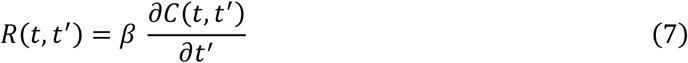

where *t*’ <= *t* and *β* = 1/*T*, *T* being the temperature of the thermal bath. In the case of non-equilibrium systems, this equality no longer holds. Instead, the linear response can be determined through generalized expressions derived from stochastic thermodynamics such as ^39^:

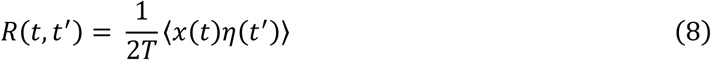

Based on this, two metrics that capture the degree of FDT violation have been introduced in prior studies ^32,38^: the differential violation *V*(*t*, *t*’) and the integral violation *I*(*t*, *t*’).

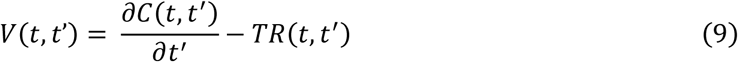

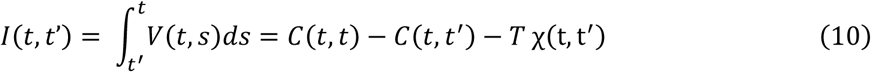

where 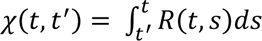 is the integrated response function, or dynamic susceptibility.

To obtain scale-free quantities that allow comparisons across datasets recorded in the same species with the same recording modality, we normalized *V*(*t*, *t*’) and *I*(*t*, *t*’) by the instantaneous autocorrelation, consistent with previous thermodynamic metrics ^31,100^.

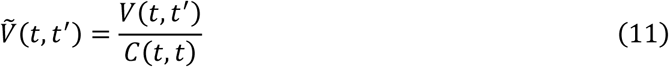

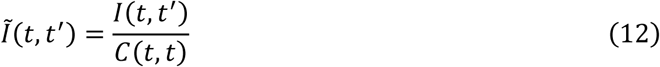

This normalization choice is justified by strict scale invariance. Normalizing by the instantaneous autocorrelation, rather than by a fixed *C*(0,0) or by *α*^2^ alone, further accounts for node-wise heterogeneity and slow variance drifts, while remaining robust to preprocessing steps that rescale amplitudes (e.g., z-scoring within runs).

Alternative estimators of FDT violations exist and are theoretically equivalent given infinite data. However, finite-sample effects can lead to variability across formulations. Following previous literature, we report the most stable estimates using the definitions of *I*^r^ as these are less prone to numerical instability ^32^.

To obtain node-level metrics of non-equilibrium, we computed the time-integrated absolute values of the violations:

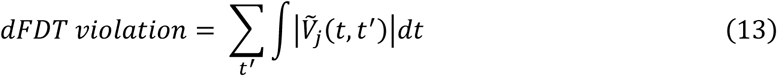

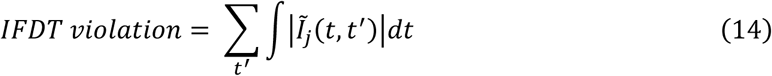

Then, for each subject, we take the average over all nodes as a more global indicator of distance from equilibrium. In the present study, we focus on these global averages. However, in conditions involving more spatially localized alterations, such as neurodegenerative or focal disorders, node-specific patterns may provide additional insight ^22^.

### Statistical Analysis

Group comparisons were conducted using non-parametric tests. For paired conditions, the Wilcoxon signed-rank test was used, followed by Holm–Bonferroni post hoc corrections. For unpaired groups, the Mann–Whitney U test was applied, and effect sizes were quantified using Cliff’s *δ*, a distribution-free, non-parametric measure directly interpretable as the probability of superiority. Binary classification performance was evaluated using the area under the curve (AROC) as a measure of discriminative accuracy.

To assess monotonic relationships between variables, Spearman’s correlation coefficients (ρ) were calculated. Pearson’s correlation coefficients (r) were also computed when relationships were assumed to be linear and variables approximately normally distributed. We tested whether the relationship between FDT violations and PCI differed across conditions using an ANCOVA that included the *FDT violations × Condition* interaction term. The significance of this interaction was further evaluated using a Freedman–Lane permutation test (10,000 permutations, within-condition shuffling) to obtain exact *p* under small-sample conditions.

In paired datasets, linear mixed-effects models (LME) were fitted to confirm the robustness of the associations between FDT violations and PCI. In mice, the LME controlled for anesthesia level (deep vs mid vs light), while in humans it controlled for anesthesia state (anesthetized vs. awake) and drug type (ketamine vs propofol vs xenon). In both cases, subject identity was included as a random intercept to account for repeated measurements.

Variables were standardized prior to regression to allow interpretation of effect sizes in standard deviation units. All p-values reported are two-tailed, and significance was defined at *p* < 0.05.

## DATA AVAILABILITY

Raw LFP recordings from eight adult male C57BL/6J mice across different anesthesia levels are publicly available at the EBRAINS repository (10.25493/WKA8-Q4T). Raw EEG recordings from neurotypical participants during wakefulness and anesthesia are publicly available at Zenodo (https://doi.org/10.5281/zenodo.806176). Additional data, including raw EEG signals from patients with disorders of consciousness, along with associated metadata, are available from the authors upon reasonable request and subject to a data sharing agreement.

## CODE AVAILABILITY

The code used for data analysis, modeling of spontaneous fluctuations, fitting the generative effective connectivity, computation of FDT violations, and other thermodynamic metrics is available at https://github.com/tomasberjagabuisan/Thermo-Consciousness.

## ACKNOWLEDGMENTS

We are grateful to all the patients and volunteers who participated in the experiments and made this work possible. We thank the members of the various laboratories involved in data acquisition for their invaluable contributions. We would like to thank Irene Acero and Paula Garcia for their insightful comments on the draft. We want to also thank Inés Blázquez for assisting with the figures.

J.M.M. acknowledges financial support from CONICET through an external scholarship for researchers. M.L.K. is supported by the Centre for Eudaimonia and Human Flourishing (funded by the Pettit and Carlsberg Foundations) and Center for Music in the Brain (funded by the Danish National Research Foundation, DNRF117). M. Corbetta is supported by Fondazione Cassa di Risparmio di Padova e Rovigo (CARIPARO) (GA 55403), the Italian Ministry of Health (NEUROCONN RF-2008-12366899, EYEMOVINSTROKE RF-2019-12369300), and the EU H2020 project euSNN, H2020-SC5–2019-2, (Grant Agreement 869505). M.V.S.V. is supported by PID2023-152918OB-I00 funded by MICIU / AEI / 10.13039/501100011033/FEDER, UE and by AGAUR 2021-SGR-01165. M. Corbetta, M.V.S.V., M.M., Y.S.P. and G.D. are supported by the project Neurological Mechanisms of Injury, and Sleep-like cellular dynamics (NEMESIS; ref. 101071900) funded by the EU ERC Synergy Horizon Europe. G.D. also acknowledges support from grant PID2022-136216NB-I00 funded by MICIU/AEI/10.13039/501100011033; by ERDF A way of making Europe, ERDF, EU; and AGAUR research support grant (2021 SGR 00917) funded by the Department of Research and Universities of the Generalitat of Catalunya.

## AUTHOR CONTRIBUTION

T.B.B., J.M.M., Y.S.P., and G.D. conceptualized and designed the study. T.B.B. and J.M.M. performed the formal analysis and developed the modeling framework. T.B.B., J.M.M., M. Cortada, M.A.C., M.V.S.V., M.M., Y.S.P. and G.D. contributed to the methodology. J.M.M, M. Cortada and S.M.G. performed complementary analysis. M. Cortada, M.A.C., M.V.S.V. and M.M. provided resources. T.B.B. performed additional data curation and filtering. T.B.B. generated all visualizations with input from M.L.K. G.D. supervised the research. T.B.B. wrote the original manuscript. All authors reviewed and edited the manuscript.

## SUPPLEMENTAL INFORMATION

Document S1. Figures S1–S8, additional methodological details and illustrations, and supplementary references.

