## Supplemental Information for "Thermodynamics of consciousness: A non-invasive perturbational framework"

### SUPPLEMENTARY INFORMATION

#### *Supplementary Note 1. Illustrative Spontaneous Traces of Electrophysiological Recordings*

To qualitatively illustrate the temporal structure of brain activity across levels of consciousness, we present representative spontaneous electrophysiological recordings from mice and humans. In mice, the LFP traces exhibit a clear progression as anesthesia deepens, with activity becoming increasingly dominated by large, stereotyped Up–Down state transitions. As isoflurane concentration increases, Up states shorten and occur less frequently, reflecting strengthened cortical bistability and the predominance of Down states (Figure S1A).

Human EEG recordings reveal an analogous state dependence. During anesthesia, the signal is dominated by large-amplitude slow oscillations, indicative of widespread cortical synchrony and reduced responsiveness. In contrast, wakefulness is characterized by lower-amplitude, desynchronized activity with faster components, consistent with active and structured neural processing (Figure S1B). Together, these examples provide qualitative insight into how spontaneous brain activity reorganizes across species as the brain transitions between conscious and anesthetized states.

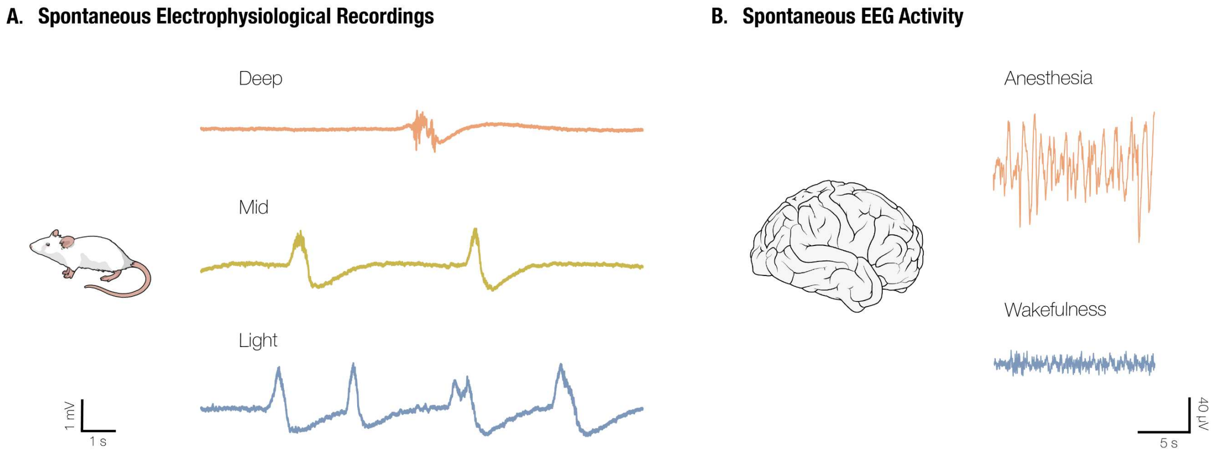


**Figure S1**. **Illustrative spontaneous electrophysiological recordings. (A)** Spontaneous LFP traces from the same mice under three different levels of anesthesia, defined by increasing isoflurane concentrations. As observed, neural activity under anesthesia becomes dominated by stereotyped Up–Down state transitions characteristic of the anesthetized cortex. **(B)** Representative human EEG patterns during anesthesia, characterized by dominant slow oscillations, and during wakefulness, marked by lower-amplitude, desynchronized activity with prominent alpha rhythms.

#### *Supplementary Note 2. FDT Violations Correlate with the Frequency of the Slow Oscillation in Animal Models*

Slow oscillations (SO) are a hallmark of low-complexity brain states. They are prominently observed during non-rapid eye movement sleep, in patients with unresponsive wakefulness syndrome, even with eyes open, and in pathological conditions such as stroke. These rhythms reflect cortical bistability, arising from alternating phases of neuronal activity and silence. Converging evidence from studies in humans, rodents, and cortical slices identifies cortical bistability and OFF periods as major disruptors of large-scale network integration and complexity.

Here, we show that the PCI correlates with the frequency of the SO (Pearson's r = 0.76, $p < 0.001$; Spearman's ρ = 0.69, $p < 0.001$; Figure S2A). Likewise, FDT violations exhibit an even stronger association with SO frequency (Pearson's r = 0.83, $p < 0.001$; Spearman's ρ = 0.80, $p < 0.001$; Figure S2B). Notably, FDT violations capture the spatiotemporal structure of SO with high sensitivity, highlighting their role as a marker of these low-complexity states.

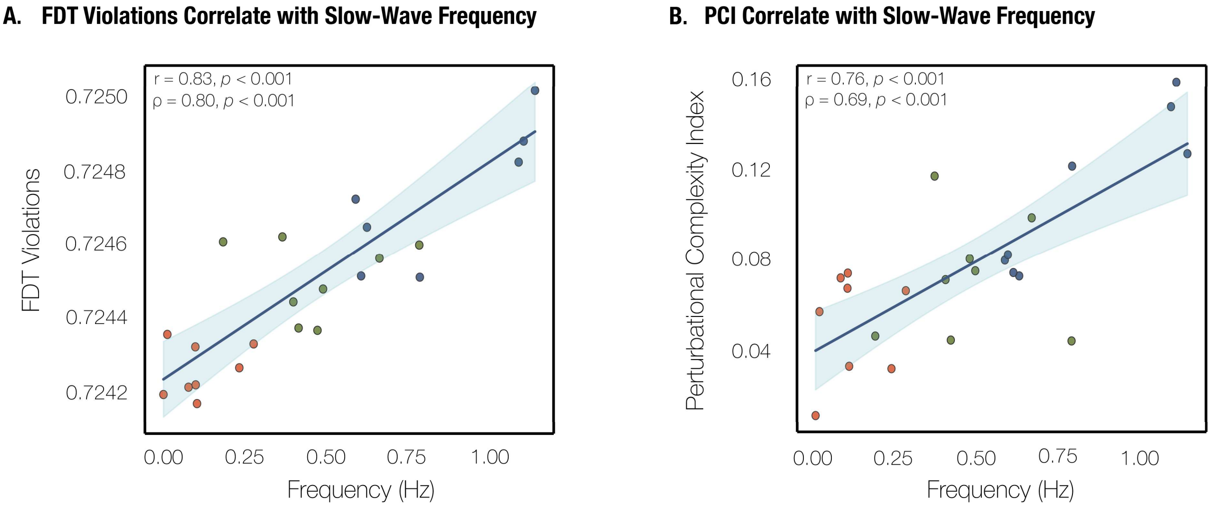


**Figure S2**. **Slow oscillation intrinsic frequency correlates with PCI and FDT violations in animal models. (A)** FDT violations show a strong correlation with the intrinsic frequency of the slow oscillations, capturing its spatiotemporal structure with high sensitivity (Pearson's r = 0.83, $p < 0.001$; Spearman's ρ = 0.80, $p < 0.001$) **(B)** PCI also correlates with the intrinsic frequency of the slow oscillation (Pearson's r = 0.76, $p < 0.001$; Spearman's ρ = 0.69, $p < 0.001$).

#### *Supplementary Note 3. FDT Violations Correlate with Other Non-equilibrium Measures*

Entropy production is one of the canonical physical measures of deviation from equilibrium. In the model used in this work, the entropy production rate (EPR) can be calculated analytically [1]. We consider a linear Langevin system described by:

$$\frac{dx}{dt} = -Bx(t) + \eta(t) \qquad (S1)$$

Where $\eta(t)$ is homogeneous white Gaussian noise $< \eta(t)\eta(t') > = 2\,\sigma^2\delta(t-t')$. From here we can determine the EPR as:

$$\phi = -\frac{1}{\sigma^2} tr(B\Theta) \qquad (S2)$$

Consider that $\Theta = BK - \sigma^2 I$, where $K$ is the stationary covariance matrix. From all the fitted brain models we calculated the EPR following Equation S2.

A key characteristic of non-equilibrium systems is their temporal irreversibility. Following previous work [2,3], we quantified this irreversibility by measuring the asymmetry of time-lagged covariances. Specifically, we computed the difference between the lagged covariance matrix and its transpose:

$$A_c = \left\| KS^{model}(\tau) - (KS^{model}(\tau))^T \right\|_{\tau=dt} \qquad (S3)$$

Where $\|.\|$ is the Frobenius matrix norm (the squared root of the squared elements), and $A_C$ quantifies the asymmetry of lagged-covariances evaluated at $\tau = dt$.

Lastly, we examined the asymmetry of the fitted coupling matrix, which has been highlighted as a key mechanism for breaking detailed balance in interacting systems [3], inducing FDT violations [4]. Following previous work [5], we characterized coupling asymmetry by means of the irreciprocity. We define the reciprocated portion of the coupling between two nodes:

$$\overleftrightarrow{B}ij = min(Bij, Bji) = \overleftrightarrow{B}ji \qquad (S4)$$

And the global reciprocity of the coupling as the ratio between the total reciprocated weight and the total connection weight:

$$r(B) = \frac{\sum_{i\neq j} \overleftrightarrow{B}ij}{\sum_{i\neq j} Bij} \qquad (S5)$$

Finally, we define the irreprocity of the coupling as $A_B = 1 - r$, which lies in the range [0,1]. A value of $A_B = 0$ corresponds to a perfectly reciprocated coupling, while $A_B = 1$ corresponds to a totally unreciprocated coupling.

We reanalyzed the three datasets incorporating the non-equilibrium measures described above.

In Figure S3 we show the results for the mice dataset. There is a reduction in EPR (Figure S3A), temporal irreversibility (Figure S3B), and coupling asymmetry (Figure S3C) with increasing anesthesia depth. These metrics significantly discriminated between light and deep anesthesia conditions (Spearman's $p < 0.05$), but they did not reliably distinguish the intermediate condition from the other two. Nevertheless, the Spearman correlation coefficients suggest that there is a track of the anesthesia state across conditions (EPR: Spearman's $\rho = 0.67$, $p < 0.001$; irreversibility: Spearman's $\rho = 0.67$, $p < 0.001$; asymmetry: Spearman's $\rho = 0.7$, $p < 0.001$). Furthermore, FDT violations increased monotonically with EPR (Spearman's $\rho = 0.72$, $p < 0.001$), irreversibility (Spearman's $\rho = 0.72$, $p < 0.001$), and coupling asymmetry (Spearman's $\rho = 0.77$, $p < 0.001$).

In Figure S4, we present the results for the three metrics in the human recordings obtained during wakefulness and anesthesia. For all participants, we observed a reduction in EPR (Figure S4A), temporal irreversibility (Figure S4B), and coupling asymmetry (Figure S4C), after xenon and propofol administration. No significant differences were observed between ketamine and wake states, consistent with our results on PCI and FDT violations.

As previously observed for FDT violations, EPR, irreversibility, and irreciprocity were significantly higher for participants who reported conscious experience (B+) compared with those who did not (B-) (Spearman's $p < 0.001$). Furthermore, FDT violations were positively correlated with EPR (Spearman's $\rho = 0.67$, $p < 0.001$), irreversibility (Spearman's $\rho = 0.67$, $p < 0.001$) and coupling asymmetry (Spearman's $\rho = 0.64$, $p < 0.001$) (Figures S4D-F).

Lastly, we analyzed the disorders of consciousness dataset (Figure S5). All non-equilibrium measures showed a decreasing tendency in patients without signs of consciousness (B-) with respect to patients with conscious signs (B+) as observed with FDT violations. However, no significant differences were found for the EPR (Figure S5A), temporal irreversibility (Figure S5B), or coupling asymmetry (Figure S5C). However, the pooled data did show a significant positive correlation between FDT violations and temporal irreversibility (Spearman's $\rho = 0.51$, $p = 0.038$).

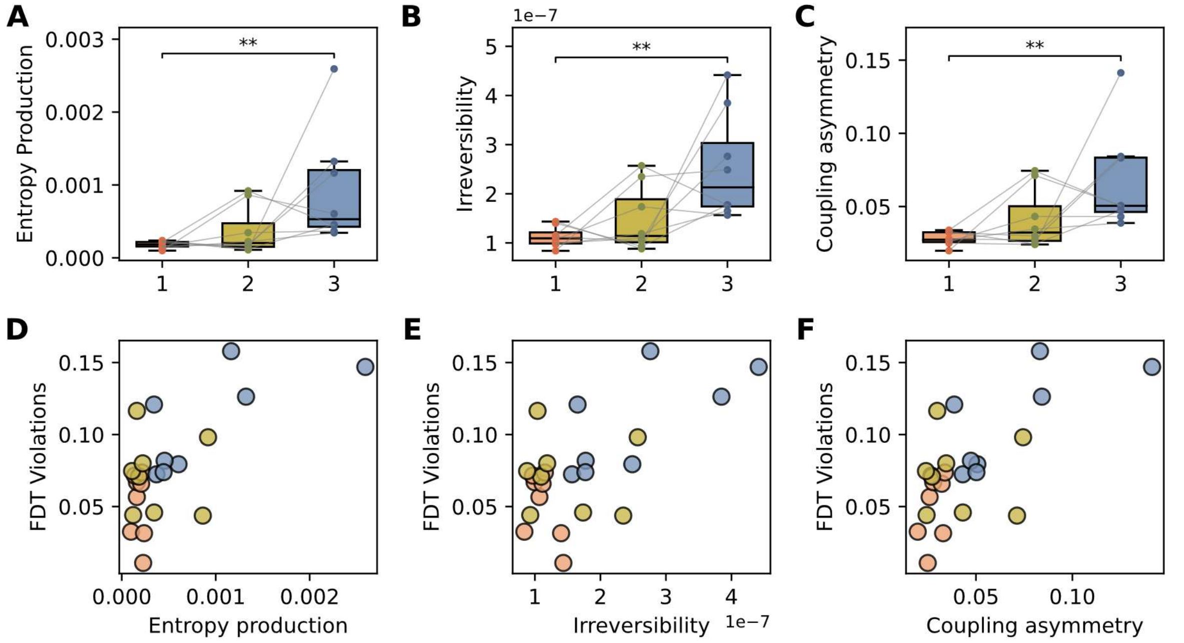


**Figure S3**. **FDT violations and different measures of non-equilibrium dynamics track levels of anesthesia in mice.** **(A)** Entropy production rate, calculated from the whole-brain model fitted to empirical spontaneous activity. (**B)** Temporal irreversibility, computed from the lagged covariances obtained from the linear stochastic model. (**C)** Asymmetry of the fitted coupling matrices. **(D)** FDT violations increased monotonically with EPR (Spearman's ρ = 0.72, $p < 0.001$). **(E)** Irreversibility (Spearman's ρ = 0.72, $p < 0.001$). **(F)** Coupling asymmetry (Spearman's ρ = 0.77, $p < 0.001$). ** $p < 0.01$

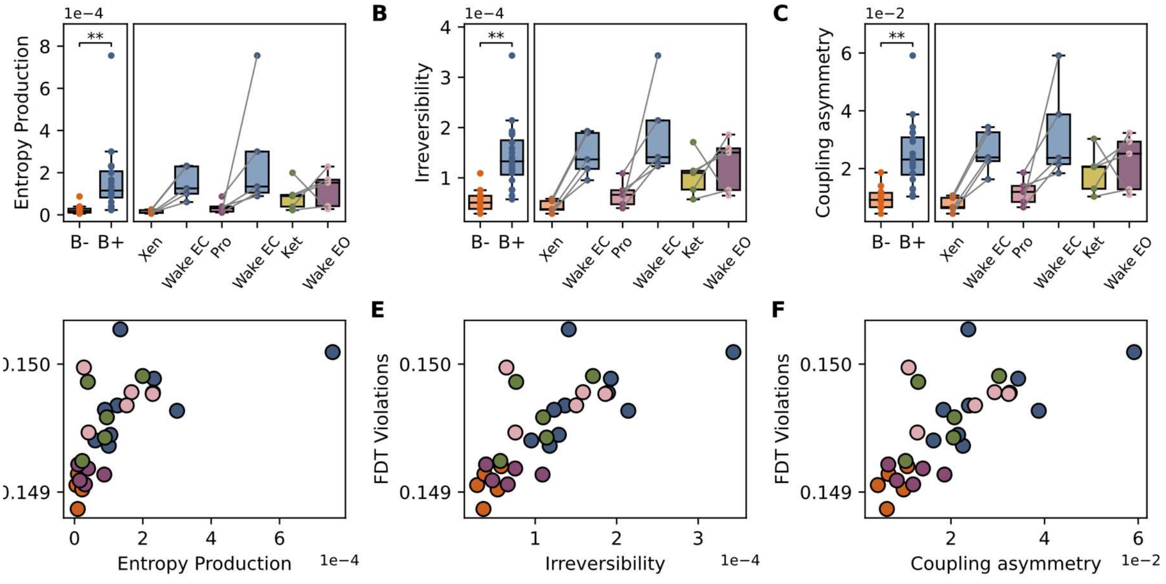


**Figure S4**. **FDT violations and different measures of non-equilibrium dynamics across levels of consciousness in wakefulness and anesthesia in humans.** **(A)** Entropy production rate obtained from the fitted model, for participants that reported conscious experience (B+) and those who did not (B-), and across anesthesia conditions. (**B)** Temporal irreversibility obtained from the linear stochastic model across anesthesia conditions. (**C)** Asymmetry of the fitted coupling across anesthesia conditions. **(D)** FDT violations increased monotonically with EPR (Spearman's ρ = 0.67, $p < 0.001$). **(E)** Irreversibility (Spearman's ρ = 0.67, $p < 0.001$). **(F)** Coupling asymmetry (Spearman's ρ = 0.64, $p < 0.001$). *** $p < 0.001$

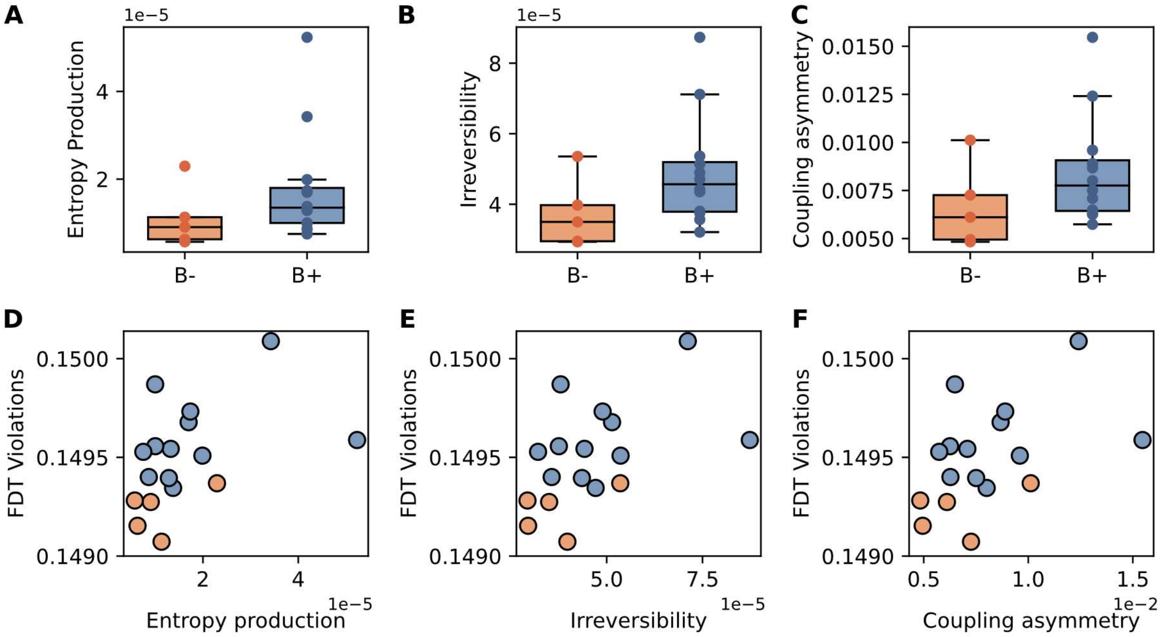


**Figure S5**. **FDT violations and different measures of non-equilibrium dynamics for conscious and unconscious patients with disorders of consciousness. (A)** Entropy production rate. (**B)** Temporal irreversibility. (**C)** Coupling asymmetry. **(D–F)** FDT violations with respect to the three non-equilibrium quantities. FDT violations increased monotonically, but a significant positive correlation was only observed between FDT violations and temporal irreversibility (Spearman's $\rho = 0.51$, $p = 0.038$)

#### *Supplementary Note 4. Extension of Human EEG Experimental Design and Data Curation*

##### *Electrode Layouts and Montages*

Figure S6 illustrates the electrode layouts and digitized montages used for the human EEG recordings in two experimental setups. Figure S6A shows the digitized electrode positions for the Nexstim EEG system (Nexstim Plc.), which consists of 60 channels and was used in a cohort of 44 participants. Figure S6B displays the digitized layout for the BrainAmp EEG system (Brain Products GmbH), comprising 64 channels and used in a smaller sample of 3 participants.

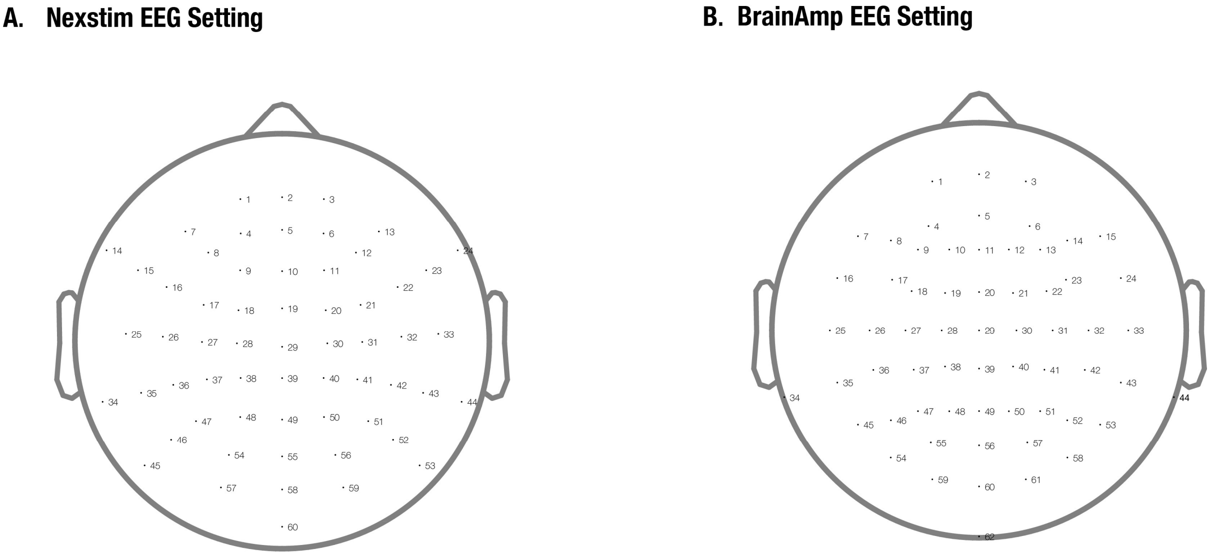


**Figure S6**. **Human EEG electrode layouts and montages. (A)** Digitization of the electrode positions for the Nexstim EEG system (Nexstim Plc., 60 channels, N = 44). (**B)** Digitization of the electrode positions for the BrainAmp EEG system (Brain Products GmbH, 64 channels, N = 3). Note the use of an external reference electrode on the forehead.

##### *Drugs and Exclusion Criteria*

Patients with disorders of consciousness (DoC) continued their standard pharmacological treatments during the EEG recording period. Specifically, antiepileptic medications were prescribed to 8 of the 17 patients. All patients in a chronic or persistent DoC state were free from sedative medications, while acute patients had been off sedatives for at least seven days prior to recording. Patients in a comatose state were excluded. Accordingly, no patient exhibited a flat or burst-suppression EEG pattern. Inclusion required at least one valid TMS-EEG session acquired either concurrently with, or in close temporal proximity to, a resting-state EEG recording.

*Independent Component Analysis for the Identification and Removal of Physiological Artefacts*

To ensure the stability of the Independent Component Analysis (ICA), we first reduced the dimensionality of the EEG data using eigenvalue decomposition, taking into account the number of rejected electrodes. Artefactual components were identified through visual inspection of each component's time series, power spectral density (PSD), and topographical distribution derived from the forward model. Ocular artefacts were typically characterized by pronounced temporal transients resembling eye movements or blinks, a PSD with elevated power at low frequencies and a steep spectral decay, and a prefrontal topography—often showing bilateral asymmetry for horizontal movements and symmetry for vertical movements and blinks. Muscular artefacts were identified by their fast, spiky temporal dynamics, occasionally organized into bursts, a broad high-frequency elevation in the PSD, and a topography indicative of localized superficial dipoles. Cardiac components were recognized by the periodic appearance of R-wave-like deflections in the time domain, a saw-tooth spectral shape associated with non-sinusoidal activity around 1 Hz, and, in some cases, a temporo-occipital distribution.

ICA-based cleaning was essential to reduce potential biases. Electrooculographic and sweat artefacts can increase low-frequency power, mimicking pathological slowing of brain activity, while electromyographic artefacts can artificially elevate power in the beta and gamma ranges, giving the false impression of increased fast cortical activity. More information can be found in the original articles of the data [6–9].

#### *Supplementary Note 5. FDT Violations in the Broadband Range in Animal Models*

To validate the focus on slow oscillations (0.1–4 Hz) in the main analysis, which account for up to 95% of the spectral power, we repeated the FDT violation and PCI correlation analyses using broadband LFP signals, within the same frequency range used for human EEG data. The results were highly consistent with those obtained in the delta range, confirming that restricting the analysis to slow oscillations preserves the relevant information while enhancing sensitivity to large-scale network asymmetries that underlie non-equilibrium dynamics.

FDT violations in the broadband range, also significantly distinguished all three levels of anesthesia (Figure S7A). This was supported by a strong monotonic (Spearman's $\rho = 0.870$, $p < 0.001$).

FDT violations correlate also with the PCI (Pearson's $r = 0.62$, $p = 0.00132$; Spearman's $\rho = 0.68$, $p < 0.001$; Figure S7B). An ANCOVA including the condition interaction term revealed no evidence of slope heterogeneity between groups, indicating that the FDT violations–PCI relationship was conserved across conditions ($F(2,18) = 0.114$, $p = 0.8933$; Freedman–Lane permutation $p_{perm} = 0.8945$).

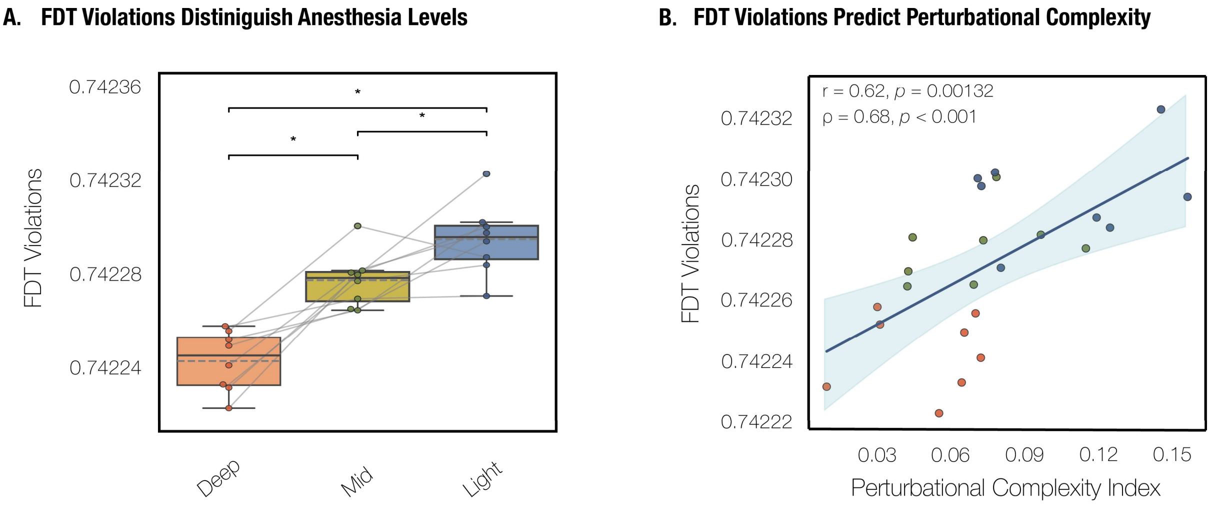


**Figure S7**. **FDT violations track levels of anesthesia in mice and correlate with the PCI in the broadband range. (A)** FDT violations calculated from the whole-brain model fitted to empirical spontaneous activity discriminated among all the conditions, as indicated by a strong monotonic relationship (Spearman's $\rho = 0.870$, $p < 0.001$). **(B)** FDT violations correlate with the PCI (Pearson's $r = 0.62$ , $p = 0.00132$; Spearman's $\rho = 0.68$, $p < 0.001$). An ANCOVA including the condition interaction term revealed no evidence of slope heterogeneity between groups, indicating that the relationship was conserved across conditions ($F(2,18) = 0.114$, $p = 0.8933$; Freedman–Lane permutation $p_{perm} = 0.8945$). All the pairwise comparisons were performed using Wilcoxon signed-rank tests and corrected post hoc with Holm-Bonferroni. * $p < 0.05$

#### *Supplementary Note 6. Illustrative TMS-EEG Traces for PCI computation*

PCI values were obtained from previous studies [6,9–11]. Here, we show two examples of TMS–EEG traces used for PCI computation (Figure S8). One example corresponds to a high PCI value, reflecting rich and differentiated evoked responses, whereas the other corresponds to a low PCI value, indicating less complex responses and a more stereotyped relationship between spontaneous and evoked activity.

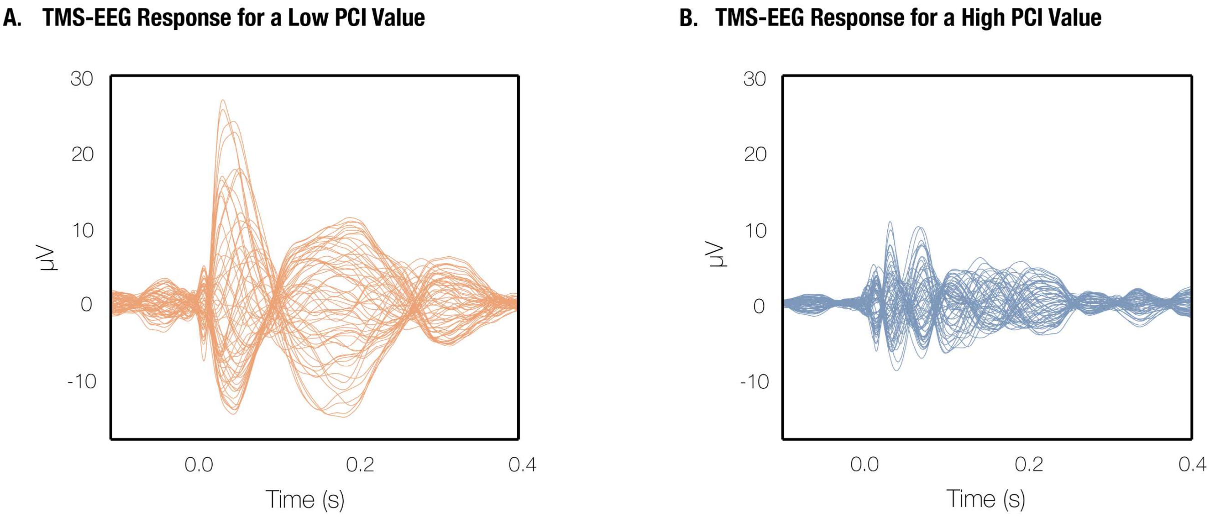


**Figure S8**. **TMS-EEG responses for PCI computation. (A)** Example of a TMS–EEG response that is less complex and more stereotyped, corresponding to a low PCI value. **(B)** Example of a TMS–EEG response that is rich and differentiated, corresponding to a high PCI value.

#### *Supplementary Note 7. Homogenous Noise in a Linear Langevin System*

In the case of homogeneous noise, the normalized stationary-state autocorrelation does not depend on the value of the noise amplitude $\sigma$ [1]. Therefore, there is no need to fit the noise, as done in previous studies, since it does not provide any additional insight [12]. In what follows, we provide proof of this.

Consider that the stationary-state autocorrelation function is given by:

$$C(\tau) = \langle x(t+\tau)x^T(t)\rangle \tag{S6}$$

In stationary conditions we can state that:

$$x(t) = \int_{-\infty}^{t} e^{-B(t-s)}\eta(s)ds \tag{S7}$$

$$x(t+\tau) = \int_{-\infty}^{t+\tau} e^{-B(t+\tau-s)}\eta(s)ds \tag{S8}$$

From $C(\tau)$, applying algebra we can derive the normalized stationary-state autocorrelation function for each node or brain region $i$.

$$\frac{C_{ii}(\tau)}{C_{ii}(0)} = \frac{[e^{-B\tau}K]_{ii}}{K_{ii}} \tag{S9}$$

In cases where the noise amplitude is identical across all brain regions or nodes $i$ (i.e. $\sigma_i = \sigma$). Then, considering $I$ to be the identity matrix:

$$Q = \sigma^2 I \tag{S10}$$

$$K = \sigma^2(B + B^T)^{-1} \tag{S11}$$

Thus, the normalized autocorrelation function for component $i$ becomes:

$$\frac{C_{ii}(\tau)}{C_{ii}(0)} = \frac{[e^{-B\tau}K]_{ii}}{K_{ii}} = \frac{[e^{-B\tau}(B + B^T)^{-1}]_{ii}}{[(B + B^T)^{-1}]_{ii}} \tag{S12}$$

From Equation S12, we can clearly see that in the case of homogeneous noise, the normalized stationary-state autocorrelation does not depend on the noise amplitude. Therefore, there is no need to fit the noise amplitude in the model, as all relevant information is embedded in the generative effective connectivity matrix $B$.
